# MCS^2^: Minimal coordinated supports for fast enumeration of minimal cut sets in metabolic networks

**DOI:** 10.1101/471250

**Authors:** Reza Miraskarshahi, Hooman Zabeti, Tamon Stephen, Leonid Chindelevitch

## Abstract

**Motivation:** Constraint-based modeling of metabolic networks helps researchers gain insight into the metabolic processes of many organisms, both prokaryotic and eukaryotic. Minimal Cut Sets (MCSs) are minimal sets of reactions whose inhibition blocks a target reaction in a metabolic network. Most approaches for finding the MCSs in constrained-based models require, either as an intermediate step or as a byproduct of the calculation, the computation of the set of elementary flux modes (EFMs), a convex basis for the valid flux vectors in the network. Recently, Ballerstein *et al.* [BvKKH11] proposed a method for computing the MCSs of a network without first computing its EFMs, by creating a dual network whose EFMs are a superset of the MCSs of the original network. However, their dual network is always larger than the original network and depends on the target reaction.

Here we propose the construction of a different dual network, which is typically smaller than the original network and is independent of the target reaction, for the same purpose. We prove the correctness of our approach, MCS^2^, and describe how it can be modified to compute the few smallest MCSs for a given target reaction.

**Results:** We compare MCS^2^ to the method of Ballerstein *et al.* and two other existing methods. We show that MCS^2^ succeeds in calculating the full set of MCSs in many models where other approaches cannot finish within a reasonable amount of time. Thus, in addition to its theoretical novelty, our approach provides a practical advantage over existing methods.

**Availability:** MCS^2^ is freely available at https://github.com/RezaMash/MCS under the GNU 3.0 license.

## 1. Introduction

Constraint-based modeling of metabolic networks has been a major subfield of systems biology thanks to its ability to identify key qualitative characteristics of networks for analyzing and extracting useful information [PRP04, BMKP14, LNP12]. A metabolic network is a collection of chemical reactions which comprise the metabolic activities (i.e., the biochemical transformation of molecules into other molecules for the purpose of maintenance and growth) of a specific organism. One important application of metabolic network analysis is to find interventions that can block a reaction of interest, typically referred to as the *target reaction*, with applications in drug target identification [HFR^+^14, TCWS06, vKK17, IB08, HBK16] and metabolic engineering [MvKK15]. When this is achieved by disabling one or more other reactions, the disabled reactions are called a *cut set*. A cut set is called “minimal” if no proper subset of it can disable the target reaction. The concept of minimal cut sets (MCS) was introduced by Klamt and Gilles [KG04] and its applications are examined in detail in [Kla06].

At the moment, the main approach used for enumerating the MCSs for a target reaction is to compute the elementary flux modes containing the target and then use a dualization procedure to produce the MCSs [GDVL17]. Here, *flux modes* are possible distributions of fluxes through the reactions, and those can be modelled as hyperedges on the vertex set of possible reactions. *Elementary flux modes* (EFMs) are flux modes which are support-minimal, and it is known that any flux mode can be written as a non-negative linear combination of EFMs. Given the full set of elementary flux modes, minimal cut sets can be obtained through the dualization of the hypergraph they define [KG04, HKS08]. Two approaches to do this are Berge’s algorithm [Ber84] and Fredman and Khachiyan’s dualization procedure [FK96]. However, both suffer from poor worst-case complexity and produce mixed results in practice. A comparatively new approach [BvKKH11] produces the MCSs without first computing the EFMs. It works by generating a dual network, which is larger than the original network and depends on the target reaction, and then computing a subset of the EFMs of that network with a specific property, which guarantees that they are precisely the MCSs in the original network. We call this the *target-specific dual network method*. In this paper, we develop a new method, MCS^2^, which also generates a dual network (either explicitly or implicitly), but in a way that is independent of the target reaction from the original network, then computes the MCSs from those EFMs of the dual network that satisfy a certain property, which also guarantees that they are precisely the MCSs for the target reaction in the original network. MCS^2^ is based on a generalization of some theoretical results by the last author [Chi10].

We implement MCS^2^ and find it to be effective on most instances we test it on. We compare it to three alternate methods for enumerating all the MCSs for a target set. The first two methods are to compute the EFMs, and then dualize them with either Berge’s algorithm, or an optimized implementation of Fredman-Khachiyan dualization, respectively. For Berge’s algorithm, we used the implementation in CellNetAnalayzer [KSRG07], containing the enhancements described in [HKS08, EMG08]. For Fredman-Khachiyan dualization, we used the recent implementation of [SSC18].

The *target-specific dual network method* [BvKKH11] first creates a dual network based on the given stoichiometric matrix and the given target reaction. It then proceeds to compute the EFMs of that dual network. Following some post-processing, the supports of these EFMs are reduced to give the required MCSs; because the MCSs correspond to only a subset of the vectors produced, this post-processing includes removing any supersets. Like MCS^2^, the target-specific dual network method reports all the MCSs without first computing the EFMs or requiring them as an input. The authors of [BvKKH11] did not provide a publicly available implementation of their method, so we did so ourselves, including all the enhancements mentioned in their supplementary materials. Most of these enhancements have improved the performance of the target-specific dual network method, for instance by reducing the size of the intermediate results.

For the majority of the models we investigated, we find that MCS^2^ is more efficient than these other methods, in terms of both running time and memory use. On the negative side, we show that our approach does not allow the enumeration of all MCSs through a given target reaction in incremental polynomial time, something that therefore remains a major open problem in the field.

Given the challenges in enumerating all MCSs (in part due to their large number, which can be exponential in the size of the network), some recent work [vKK14, VMRR16] uses Mixed Integer Linear Programming (MILP) formulations to enumerate a subset of the MCSs, in increasing order of size. In some practical applications, quickly obtaining a few MCSs of minimum size may be more desirable than enumerating the complete set. We therefore adapt the MCS^2^ approach to use MILP formulations to address this task. We also implement this method, which we call MCS^2^-MILP, and compare it to MCSEnumerator, the target-specific dual network approach adapted to MILP [vKK14]. The comparison shows that MCS^2^-MILP performs at least as well as MCSEnumerator using a state-of-the-art MILP solver [IBM].

We conclude that MCS^2^ is a promising approach for the computation of MCSs in metabolic networks, and expect it to be a beneficial addition to the analysis tools available for metabolic network models.

### 1.1 Definitions

We now introduce the terminology we will be using throughout this paper. When we speak of a metabolic network, it is understood that we are talking about a model in the constraint-based modeling formalism.

#### Definition 1 (Stoichiometric matrix)

The *stoichiometric matrix S* is an *m × n* matrix with each row representing a metabolite (indexed from 1 to *m*) and each column, a reaction (indexed from 1 to *n*). The entry *S*_*ij*_ indicates how many units of metabolite *i* are produced (if *S*_*ij*_ > 0) or consumed (if *S*_*ij*_ < 0) by reaction *j*. A vector *v* is *feasible* with respect to *S* if it is in the *null space* of *S*, i.e., if it satisfies *Sv* = 0.

#### Definition 2 (Reaction irreversibility)

The set 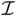 of *irreversible* reactions is a subset of the set of reactions constrained to have only non-negative fluxes. Its complement 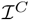, the set of *reversible* reactions, is allowed to have fluxes of any sign. A vector *v* respects the reaction irreversibility constraints if 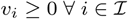, also written as 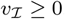.

#### Definition 3 (Metabolic network)

A metabolic network 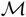 is a pair 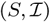, where 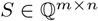 is a stoichiometric matrix and 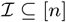 is the set of irreversible reactions. A vector *v* is a *flux mode* if it is feasible with respect to *S* and respects the irreversibility constraints, i.e., *Sv* = 0 and 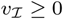. The set of all such vectors is called the network’s *flux cone*.

#### Definition 4 (Reconfigured network)

Let 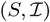 be a metabolic network. We can *reconfigure* this network by replacing *S* with 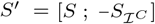 and 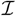 with *∅*. This is equivalent to splitting each reversible reaction in the network into its forward reaction and reverse reaction.

#### Definition 5 (Null space matrix and network)

Let *S* be a matrix. A *null space matrix* of *S* is a matrix whose rows form a basis of the null space of *S*. The *null space network* of a metabolic network with stoichiometric matrix *S* is the fully reversible metabolic network (i.e., with 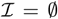) whose stoichiometric matrix is a null space matrix of *S*.

#### Definition 6 (Positive and negative support)

Let *v* be a vector. The *positive support* of *v*, 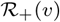, is the set of positions *i* where *v*_*i*_ is positive: 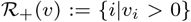. The *negative support* of *v*, 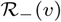, is the set of positions *i* where *v*_*i*_ is negative: 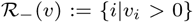. Their union 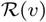 is the *support* of 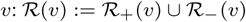.

#### Definition 7 (Coordinated support)

Let *v* be a vector of size *n* and let *A ⊆* [*n*] be a set of coordinates. The *A-coordinated support* of *v*, 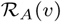, is the union of its negative support on the positions in *A* and its support everywhere else: 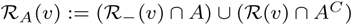.

#### Definition 8 (Elementary flux mode)

Let 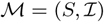 be a metabolic network, and let *v* be a flux mode of 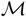. It is an *elementary flux mode* (EFM) if its support is minimal among all the flux modes of 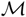, i.e., 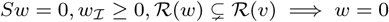 [SFD00, GK04].

#### Definition 9 (Minimal cut set)

Let 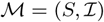 be a metabolic network, and let *t* be a reaction. *C* is a *cut set* for *t* if *Sv* = 0, 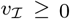, *v*_*C*_ = 0 = ⇒ *v*_*t*_ = 0. *C* is a *minimal cut set* (MCS) if it is inclusion-minimal: *D* ⊊ *C* = ⇒ ∃*v* s.t. *Sv* = 0, 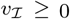, *v*_*D*_ = 0, *v*_*t*_≠ 0 [KG04].

#### Definition 10 (Canonical form of a network)

Let 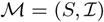 be a metabolic network. We say that 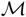 is in *canonical form* if it satisfies:

1. No blocked reactions: for every reaction *i*, there exists a flux vector *v* with *v*_*i*_ = 1;
2. Proper directedness: for every reaction 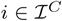, there exists a flux vector *w* with *w*_*i*_ = −1;
3. No enzyme subsets: no reaction pair *i* ≠ *j* satisfies *v*_*i*_ = *κv*_*j*_ with 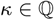 for all flux vectors *v*;
4. No redundant constraints: *S* has full row rank.

A metabolic network can be reduced to an equivalent one in canonical form (a.k.a. compressed form) in time polynomial in *m* and *n* [Chi10].

## 2 The MCS^2^ method

Let *S*_*i*_ be the *i*-th row of the stoichiometric matrix *S*. Then *S*_*ir*_ represents the amount of metabolite *i* consumed or produced by reaction *r* (in these cases, *S*_*ir*_ < 0 and *S*_*ir*_ > 0, respectively). Assume that reaction *r* produces metabolite *i* if it has a positive flux. Then, in a steady state where no reaction consuming metabolite *i* is active, reaction *r* must be inactive in the forward direction. If reaction *r* happens to be reversible, it must be consuming metabolite *i*, and its flux must be negative. This shows that reaction *r* is blocked in the forward direction if we disable every reaction that can consume metabolite *i*, i.e., every irreversible reaction with a negative value in row *i* and every reversible reaction with a non-zero value in row *i*. The set of such reactions is then a cut set for the forward direction of reaction *r*. Every row gives us some, not necessarily minimal, cut set in this manner.

We can apply the same reasoning to linear combinations of the metabolites. Consider a new *virtual* metabolite *x*, which represents a linear combination of rows *S*_*i*_ and *S*_*j*_ corresponding to metabolites *i* and *j* respectively, say for example *u*_*x*_ = 2*S*_*i*_ − *S*_*j*_.

Since the fluxes producing and consuming each metabolite are balanced in any admissible vector, so are the fluxes of their linear combinations, so the virtual metabolite *x* must also be balanced. If we pick a reaction with a positive value in *u*_*x*_, it produces a virtual metabolite *x* when it has a positive flux. It will therefore be blocked if we cut all irreversible reactions with negative values in *v*_*x*_ and all reversible reactions with non-zero values in *v*_*x*_. Thus, we can obtain cut sets from the vector *u*_*x*_, which is a member of the row space of *S*, as we did with *S*_*i*_ and *S*_*j*_. A proposal for finding cut sets by analyzing the row space of the stoichiometric matrix was introduced in the Ph.D. thesis of Chindelevitch [Chi10]. The intuition we described shows how vectors in the row space can generate cut sets. However, the lemmas proven in [Chi10] only work for the fully reversible networks (i.e., with 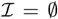) or fully reversible networks (i.e., with 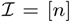). We generalize them here to networks with both irreversible reactions and reversible reactions.

### 2.1 Enumerating the full set of MCSs

In the MCS^2^ method, the dual network is the null space network of the original network, i.e., a fully reversibile network whose stoichiometric matrix is the null space matrix of the original stoichiometric matrix. The EFMs in this dual network map to minimal cut sets of the original network, though, as we will see, the mapping can be many to one. The dual network has the same number of reactions, *n*, as the original one, but it typically has fewer metabolites; if the original network has *m* metabolites and the stoichiometric matrix is full rank, the dual has *n* − *m* metabolites.

#### Lemma 1 (MCSs for an irreversible reaction)

Let 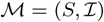 be a metabolic network. Let 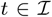 be an irreversible target reaction. Then *C* is a cut set for *t* if and only if there exists a vector *u* in the row space of *S*, *u* ∈ *Row*(*S*), such that *u*_*t*_ = 1 and 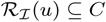.

Proof. This lemma is an extension of Lemma 3 of [Chi10]. We observe that *C* being a cut set for irreversible reaction *t* is equivalent to:

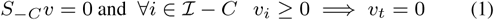

Based on Farkas’ Lemma and the irreversibility of *t*, we only need to find a constraint that implies *v*_*t*_ ≤ 0. Thus, there exists a *y* such that:

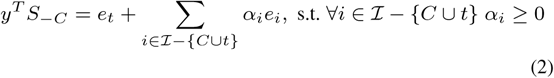

Here, *e*_*i*_ is a vector with a 1 in the *i*-th position and 0 elsewhere. Thus:

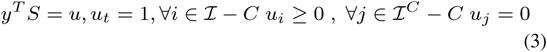

Therefore 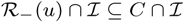 and 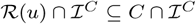, and so 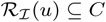.

For the other direction, suppose that *u* satisfies equation (3) for some *C*. Then the union of the indices *i* in 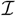 for which *u*_*i*_ < 0 and the indices *i* in 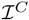 for which *u*_*i*_ ≠ 0 is a subset of *C*, i.e., 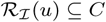. Then equality (2) holds and, by Farkas’ lemma, so does condition (1).

#### Lemma 2 MCSs for one direction of a reversible reaction)

Let 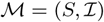 be a metabolic network. Let *t* be a reversible target reaction. Then *C* is a cut set for the forward (reverse) direction of *t* if and only if there exists a vector *u* ∈ *Row*(*S*) such that 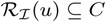 and *u*_*t*_ = 1 (*u*_*t*_ = − 1), respectively.

Proof. If we assume that *t* is irreversible for a moment, the first part already follows from the previous lemma. For the second part, replace *t* with −*t* in *S* to create *S′*. Reaction *t* is blocked in the forward direction in *S′* if and only if reaction *t* is blocked in the reverse direction in *S*, and there is a bijection between the vectors in *Row*(*S*) and those in *Row*(*S′*) via the mapping that multiplies the *t*-th coordinate by *−*1.

With these lemmas, Algorithm 1 below can be used to find the minimal cut sets for a set of target reactions *T* = {*t*_1_, *t*_2_, …, *t*_*k*_} in an arbitrary metabolic network 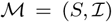, where *T* has separate elements for the opposite directions of a reversible reaction. We call this method MCS^2^ because it computes MCSs as the minimal coordinated supports (MCSs) of the elementary flux modes in the dual network.

The MCS^2^ method computes the null space network of the original network. Then, by applying coordinated support to the EFMs in which each target reaction is active in turn, it generates the cut sets related for this target. All the MCSs are among these cut sets, and are obtained by pruning. The null space network is fully reversible, and since the null space of the null space is the original space, the dual network of the dual network is equivalent to the original network all of whose reactions have become reversible. An example is shown in Figure 1, where the dual network appears alongside the original one. As shown in the example, flux modes with an active target reaction in the dual network map to cut sets for the target reaction in the original network.

**Figure 1:**
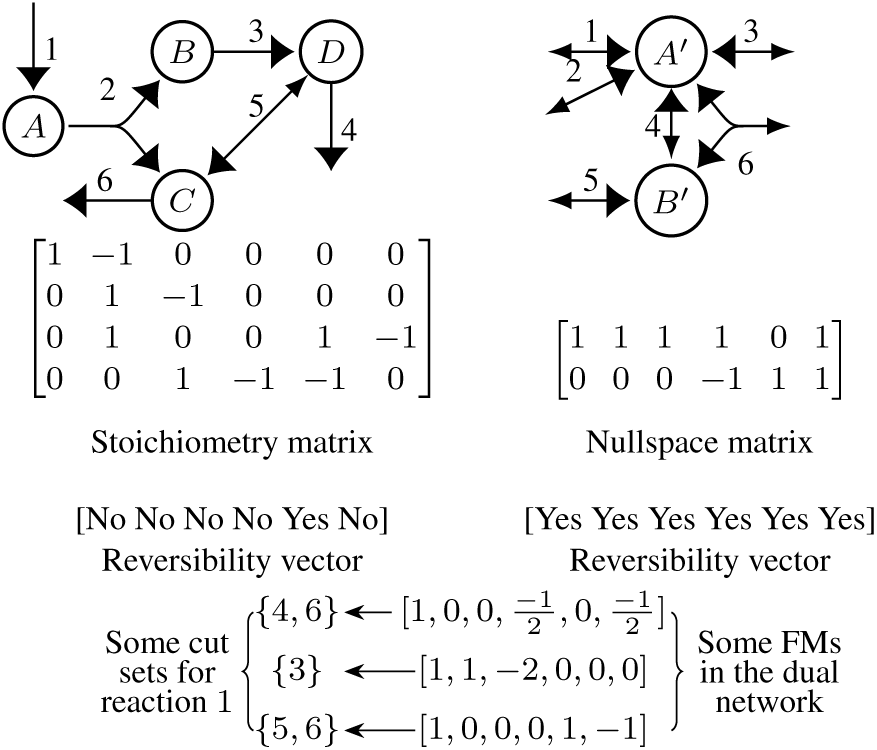
Example of a metabolic network with its associated dual network. Some of its FMs involving target reaction 1 are shown; their 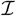-coordinated supports result in cut sets for it in the original network.

#### Algorithm 1 MCS enumeration via the MCS^2^ method

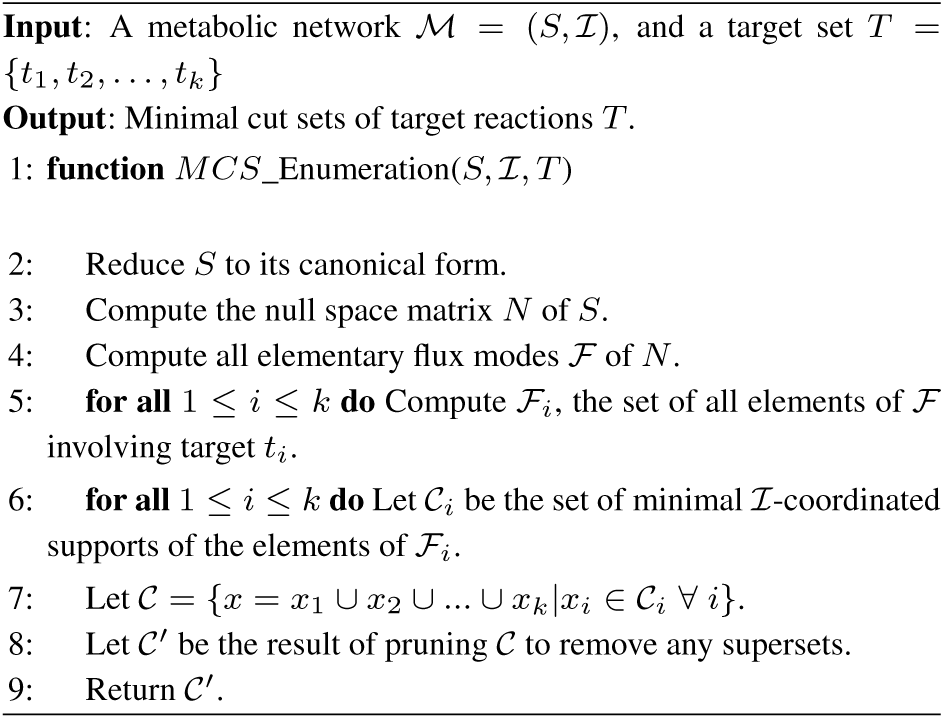

Flux modes finders such as FluxModeCalculator [vKWvD15] reconfigure the network before applying the double description method. The double description method is an algorithmic approach for finding the extreme rays of a pointed cone described by linear constraints. The reconfigured network is *N′* = [*N*; *−N*], where *N* is the null space network of *S*. Figure 3 shows the null space matrix of *N′*, which is the starting point of double description method. The double description method begins by using elementary row operations to put the matrix in the form suggested in [Wag04] which contains an identity matrix of size *m* + *n*. When the double description method finishes, it outputs the extreme rays describing a cone in 2*n*-dimensional space [TS08]. These extreme rays are the non-zero vectors in the flux cone with minimal support. On the other hand, 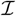-coordinated support does not count non-zero values in some dimensions, namely, those that correspond to positive values in irreversible reactions. If we ignore these dimensions, we project the cone into a lower-dimensional subspace. While the image of a pointed cone remains a pointed cone, the extreme rays of the new cone are those in the original flux cone with minimal support in the remaining dimensions. Figure 2 shows why all the minimal coordinated supports are among the minimal supports, while arguing that there may be some redundant results among them as well. The next theorem formalizes this:

**Figure 2:**
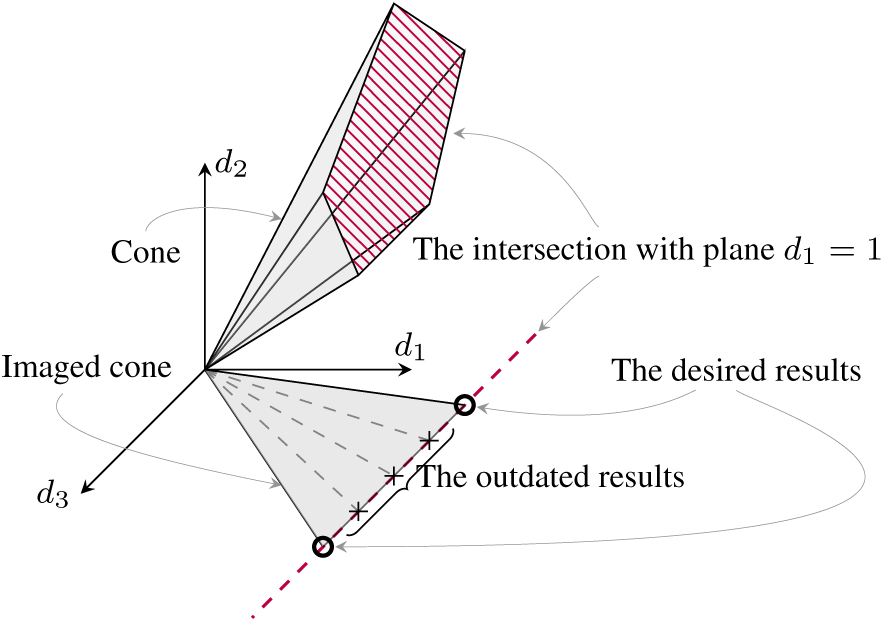
Each extreme ray of the projected cone is an image of an extreme ray in the original cone, while some extreme rays of the original cone do not project to extreme rays. It is also possible that two or more extreme rays in the original cone project onto the same one. Our desired projections lie in the plane where the value in the target position is 1.

**Figure 3:**
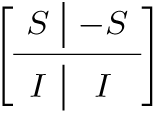
This (*m* + *n*) × 2*n* matrix is the nullspace of the reconfigured nullspace of stoichiometry matrix *S*. The double description method begins on this space and finds extreme rays with length 2*n*.

#### Theorem 1 (Correctness of the method)

Algorithm 1 returns precisely the set of minimal cut sets of the network 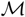 for a single target reaction.

Proof. Let *t* be the target reaction. We prove the inclusion in both directions. First, let 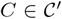 be one of the sets returned by the method above. Then *C* is a cut set for *t* in the reconfigured network, by Lemma 1 and by construction. Indeed, 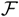 contains flux modes of *N* involving *t*, which are precisely the vectors in the row space of *S* involving *t*, and 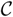 (as well as 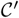) contains the 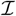-coordinated supports of these vectors.

Now, let *C* be a minimal cut set for *t* in 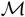. We will show that 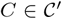. By lemma 1, there exists a vector *u* ∈ *Row*(*S*) such that *u*_*t*_ = 1 and 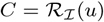. Since *u* ∈ *Row*(*S*) ⇐⇒ *u* ∈ *Null*(*N*), *u* is a conical combination of the elementary flux modes of *N*. Note that the results of Müller and Regensburger [MR16] imply that since the space to which *u* belongs is linear (i.e., it does not need to satisfy any non-negativity constraints), this conical combination can be chosen to be **conformal**, meaning that there are no cancellations involved in any component. Let such a conformal conical combination be given by

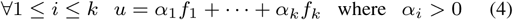

Since all the coefficients are strictly positive in (4), we deduce that

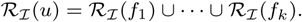

Indeed, each 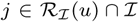 must have a negative component in at least one of the *f*_*i*_, as otherwise the *j*-th component of the right-hand side of (4) will be non-negative, which gives the ⊆ direction, and the fact that the combination is conformal gives the ⊇ direction, as otherwise there would be a cancellation.

In particular, we deduce that 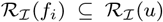 for each 1 *≤ i ≤ k*. In this case, the minimality of *C* implies that either 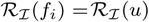 or *f*_*i*_ has a 0 in position *t*, for each 1 *≤ i ≤ k*. But since *u* has a 1 in position *t*, there must be at least one *f*_*i*_ in the first category, so that 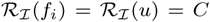 and therefore, 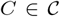. Once again, by the minimality of *C* we conclude that 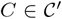 since it cannot be a superset of the 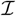-coordinated support of another vector in 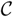, concluding the proof.

#### 2.1.1 Limitations

Our method is limited to blocking one direction of a given reaction. However, in practice, blocking one direction of a given reaction is the typical objective [LB07]. To block multiple reactions it is possible to compute the MCSs of every target reaction, take unions of all possible combinations, then remove the supersets. However, this may not always be efficient.

A more critical issue is the possibility of generating a large number of non-minimal cut sets before the post-processing. The following Lemma shows that this type of blow-up can occur in theory:

##### Lemma 3 (Large number of supersets in the final step)

For every integer *k ≥* 2 there exists a network containing *k* + 2 metabolites, 3*k* +3 reactions and 2^*k−*1^ +1 elementary vectors for the target reaction *t* = 1 that map to the exact same minimal cut set. This network is in canonical form, and is elementally balanced as per [ZSBC18].

Proof. We construct the network as follows. There are two special metabolites, denoted *M*_*I*_ (initial) and *M*_*F*_ (final), and *k* intermediate metabolites, denoted *M*_*i*_ for 1 *≤ i ≤ k*. For each metabolite, we have an export reaction and an import reaction, with the export reactions for each intermediate metabolite coupled with an import of the final metabolite. Lastly, each intermediate metabolite except the first one can be transformed into the first one, *M*_1_, which itself can also be transformed into the initial metabolite *M*_*I*_. All reactions in the network are irreversible and all the stoichiometric coefficients are *±*1.

We order these reactions as follows (for simplicity of argument):

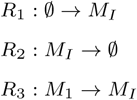

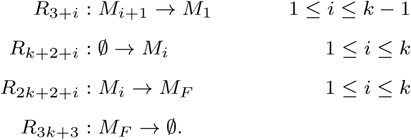

The stoichiometric matrix then looks as follows (shown for *k* = 2):

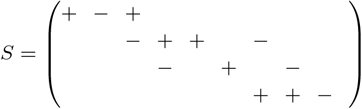

Here, a + represents a 1 and a *−* represents a *−*1. We now proceed to show each part of the desired statement:

- The network is elementally balanced because every reaction that is not pure import or pure export is an exchange of one metabolite for another in a 1-1 ratio, so we can consider each metabolite as containing exactly 1 atom.
- The network is in canonical form because every reaction can be active and no pair of reactions is constrained to have proportional fluxes; this is evidenced by the following flux modes:

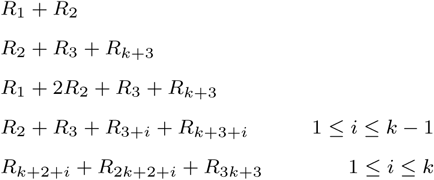 This set of fluxes includes every reaction at least twice, and in at least two of these the sets of other active reactions are disjoint. The only exceptions are *R*_1_ and *R*_3_, which need *R*_2_ to be active in order to occur, but the first and third flux modes (respectively second and third flux modes) show that their fluxes are not proportional to that of *R*_2_ or to each other; and and the reactions *R*_3+*i*_ and *R*_2*k*+2+*i*_ for 1 *≤ i ≤ k −* 1, both of which need *R*_*k*+3+*i*_ to be active in order to occur, but not in a fixed ratio, as evidenced by taking linear combinations of the last two sets of flux vectors.
- The stoichiometric matrix has full row rank, i.e., no metabolite generates a redundant constraint, because every metabolite except *M*_*F*_ has a pure import reaction, while *M*_*F*_ has a pure export reaction.
- There is a unique MCS for target reaction *R*_1_, namely, *R*_2_. This is because *R*_2_ is the only reaction consuming *M*_*I*_ (recall that all reactions are irreversible). The first row of *S*, *u* := *e*_1_ *− e*_2_ + *e*_3_ produces this MCS via its negative support.
- Lastly, there are 2^*k−*1^ additional vectors in the row space of *S* that produce supersets of this MCS via their negative support. The first one is obtained by adding the second row of *S* to *u*, replacing it by

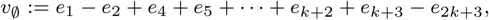

and then picking any subset *P* of the set of *k* − 1 entries *E* := {4, 5, …, *k* + 2} to form a new vector *v*_*P*_, as follows. Let 3 + *j ∈ P* be an element of the chosen subset, where 1 *≤ j ≤ k −* 1. We will replace *e*_3+*j*_ with *e*_*k*+3+*j*_ − *e*_2*k*+3+*j*_ via the addition of the (*j* + 2)-nd row of *S* (corresponding to the intermediate metabolite *M*_*j*+1_) to the starting vector. Indeed, this row contains three non-zero entries: a *−*1 from reaction *R*_3+*j*_ (which cancels out the 1 in position *e*_3+*j*_), as well as another 1 from reaction *R*_*k*+3+*j*_ and another *−*1 from reaction *R*_2*k*+3+*j*_. We do this addition independently for each element of *P* to get *v*_*P*_ (if *P* = *∅* we get *v*_∅_). It is easy to check that *v*_*P*_ has support:

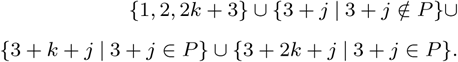 No proper subset of this support can produce a non-trivial vector in the row space of *S*, as it is impossible by construction to add a linear combination (possibly with negative coefficients) of the rows of *S* to *v*_*P*_ without adding any new elements to its support, so each *v*_*P*_ is elementary. Furthermore, the negative support of *v*_*P*_ is:

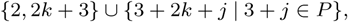

which is a strict superset of the negative support {2} of *u*.

#### 2.1.2 Advantages

An advantage of the MCS^2^ approach is that we find the MCSs directly, without first computing the EFMs of the original network. Also, we do not need to reconfigure or alter the stochiometric matrix; every step is performed directly on the original stoichiometric matrix or its null space matrix. Network compression or reduction may be done as a preprocessing step before going through the main procedure, but these are only used to reduce the running time and space and are optional. These advantages are shared with the target-specific dual network approach.

However, there are additional advantages that MCS^2^ has over this method. First, the null space matrix is typically smaller than the original matrix, especially if the original matrix is nearly full-rank, while the target-specific dual network has a matrix that is always larger. This difference in input size can lead to substantial resource savings during EFM computation. Second, and perhaps most importantly, the dual network is independent of the target reaction in our method, while it is not with the target-specific approach. This means that we can calculate these EFMs once and use them for any given target reaction to be blocked.

### 2.2 Generating small MCSs

An alternative strategy for computing EFMs is via mixed integer linear programming (MILP), particularly when only a few small MCSs are required instead of a full enumeration [RPdF^+^13, RPT^+^14]. Recall that EFMs are minimal-support vectors in the null space. Our method for finding small MCSs, which we call MCS^2^-MILP, similarly looks for vectors with minimal coordinated support in the row space, which is to say, EFMs with minimal coordinated support in the dual network.

#### Lemma 4

(MCSs of a target set of reactions in a fully irreversible metabolic network). Let *S* be the stoichiometric matrix of a fully irreversible metabolic network 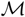. Let *T* be a set of target reactions. Then *C* is a cut set for all the reactions in *T* if and only if there exist a vector *u ∈ Row*(*S*) such that 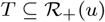 and 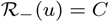.

Proof. We need to show that every cut set for the target reactions arises from a vector in the row space with the described constraints, and every vector in the row space satisfying those constraints maps to a cut set.

Let *C* be a cut set for all reactions in *T*. Therefore, *C* is a cut set for each reaction in *T* = {*t*_1_, *t*_2_, …, *t*_*k*_} individually. Based on Lemma 1, there exist vectors *u*_1_, *u*_2_, …, *u*_*k*_ ∈ *Row*(*S*) such that 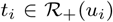 and 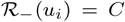 for 1 *≤ i ≤ k*. In other words, for all vectors *u*_*i*_ (1 *≤ i ≤ k*) the only negative elements are the ones with indices belonging to *C*, and all other elements are non-negative, with a strictly positive value in the one with index *t*_*i*_ in the vector *u*_*i*_, for 1 *≤ i ≤ k*. If we define the vector *u* := *u*_1_ + *u*_2_ + *…* + *u*_*k*_, then 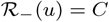 and 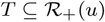, and *u* is clearly in *Row*(*S*).

Now, let *u* be a vector in *Row*(*S*) such that 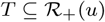 and 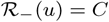. Then 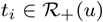 for all 1 *≤ i ≤ k*. Based on Lemma 1, 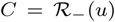 is a cut set for the reaction *t*_*i*_, for each 1 *≤ i ≤ k*. Therefore, *C* is a cut set for all the reactions in *T*, completing the proof.

Based on this Lemma we are able to find minimal cut sets for every set of target reactions without the restriction of only blocking one direction of a reaction. Since reversible reactions can be split into two reactions after reconfiguration, we can block a reversible reaction in one direction or in both directions.

Let *S′* be the *m × n′* reconfigured matrix of the *m × n* stoichiometric matrix *S* with irreversible reactions 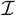. Since all the values in the stoichiometric matrix are proportions of consumed and produced metabolites, we can scale each row of *S* to have only integer entries without changing its structural properties.

We now describe how to encode the problem of finding the smallest MCS for a target set *T* as a mixed-integer linear program (MILP).

Let 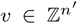 be a vector in the row space of the reconfigured matrix *S′* corresponding to the smallest MCS for target reaction set *T* = {*t*_1_, *t*_2_, …, *t*_*k*_}. Then there exists a vector 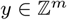 s.t *y*^*T*^ *S* = *v*. If we define *r*^+^, *r*^−^ ∈ {0, 1}^*n*′^ as the indicator vectors of the positive and negative supports of *v*, respectively, we may force *v*_*i*_ to be non-positive if 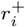 is 0, and force it to be non-negative if 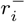 is 0, by adding the following constraints using 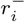 and 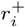 as indicator variables:

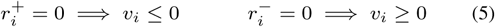

There must also be positive values in the target positions:

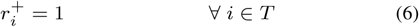

These constraints also ensure that *v* = 0 is not in our feasible space. To make *v* a vector in the row space of *S′* we need to add the *y*_*j*_ variables, namely, the entries of a vector *y* with size *m*. The constraint *y*^*T*^ *S′* = *v* then ensures that *v* is an element of the row space of *S′*.

The objective function for finding the smallest minimal cut set is:

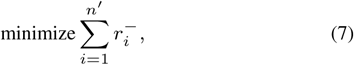

since the cut set is precisely the negative support of *v*, i.e *r*^−^.

Suppose that we have found the smallest MCS 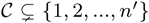. To find the next smallest MCS, we need to exclude 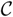 and all its supersets from our feasible space. This is achieved by the following constraint:

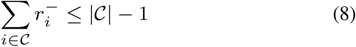

We can keep excluding newly found MCSs and thus enumerate them in order of increasing size. As we stated above, in most scenarios we only wish to block an irreversible reaction or one direction of a reversible reaction. In those cases, we can avoid re-configuring the network to have a smaller stoichiometric matrix. Let *t* be the only target reaction. Instead of the constraints (6), we only need one constraint 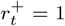 if we want to block it in forward direction, and we need the constraint 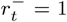 if we need to block it in the reverse direction. The objective function (7) and constraints (8) can be updated as follows to reflect the coordinated support instead of the negative support:

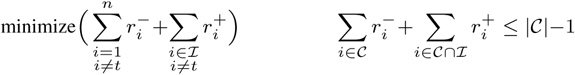

Unlike [vKK14], our problem formulation does not require any additional constraints, because they only reduce a part of the feasible space of our problem without affecting the optimum objective value. This concerns constraints such as 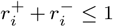 and 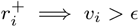.

## 3 Implementation details

Except where noted, the implementations we discuss are in MATLAB. Each method that we consider requires an extreme ray computation, with the underlying cone varying. We used FluxModeCalculator’s EFM generator [vKWvD15] for this purpose. Note that the optimized Berge algorithm implemented by CellNetAnalyzer [KSRG07] uses the older EFM finder of CellNetAnalyzer by default. However, we observed that it is a slower implementation of an identical calculation, so we rewrote this part to use FluxModeCalculator in order to make a fair comparison. The MCS^2^ method and the target-specific dual network method both need to remove redundant supersets from the obtained extreme rays, since the desired minimality is not with respect to the full support of the vector, but only a specific part of it (the 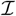-coordinated support for the former, and support in the *v*-coordinates for the latter). We use an implementation in Java whose time complexity is *O*(*N* ^2^) for a collection of *N* sets. All stoichiometric matrices are compressed by the Mongoose [CTRB14] before processing, which converts them to a canonical form.

Since the null space is needed for the row space method, we calculate the null space basis matrix using Mongoose [CTRB14]. Since finding the MCSs in every method takes several seconds to several minutes, and the computation time of the null space basis matrix is less than a second in every case, we ignore this component of it. The reduced matrix given by Mongoose (for Berge and MFK), the target-specific dual matrix (for the target-specific dual method), and the null space basis matrix (for MCS^2^) get further compressed by FluxModeCalculator before processing. For the Berge algorithm we used CellNetAnalyzer [KSRG07]. We also used an existing implementation of the improved modified Fredman-Khachiyan (MFK) algorithm [SSC18]. However, we implemented the target-specific dual method from scratch using MATLAB and the source code of FluxModeCalculator. All the enhancements mentioned in the supplementary material of the original paper [BvKKH11] were implemented as well.

We used CPLEX [IBM] to solve the MILPs. The implementation was done via the Java API and has been implemented for single target reactions without network reconfiguration, and for multiple target reactions with network reconfiguration. Since the stoichiometric matrices needed to contain only integers, we used the `integralize` function of MONGOOSE [CTRB14] to multiply each row by the smallest possible integer that makes all the values integer (which is the least common multiple of the denominators of its entries). We also tested the results of our MILP in small networks against other implementations to make sure that the results are consistent. The implementation of all the methods and the MILP version of our method are publicly available at https://github.com/RezaMash/MCS under the GNU 3.0 license. Some of them require the use of non-public modules available for academic use, such as CellNetAnalyzer [KSRG07] and CPLEX [IBM].

## 4 Results

In this section we summarize the performance of MCS^2^ and MCS^2^-MILP in comparison to the other methods.

We ran the implementations on the first five models in our database in the GitHub repository. We compared the set of MCSs in these five small examples to confirm that all the implementations produced same results. For the other results presented in the tables we again checked the number of MCSs reported by implementations if they finished, and the numbers matched in all cases. We then ran the methods on several models from the BioModels database [LDR^+^10]. There were a few models on which our method either was not able to finish in the given time (5 hours) or took much longer to report the MCSs, while the optimized Berge was able to finish in time and beat our method (see Table 3 for an example). This is due to the large number of supersets generated in that example by MCS^2^. However, MCS^2^ always performed better than the target-specific dual network approach, despite all its suggested enhancements being implemented. In addition, as can be seen for the *hepatic polyamine and sulfur aminoacid combined* model [RPMSJM12], the Berge and MFK methods could not finish in five hours, but MCS^2^ generated results in 4 minutes, and the dual method in 30 minutes.

The first task of every method is an extreme ray computation, which for Berge and MFK is the well-known EFM computation. Berge and MFK then proceed to generate the MCSs through dualization, while the secondary process of the target-specific dual network and MCS^2^ approaches is removing the redundant cut sets. In the first two provided examples, the target is the forward direction of the first reaction. Table 2 shows the computation time for calculating the MCSs for all possible target reactions. In the *kinetic model of yeast metabolic network*, described in [SLS^+^13], our method’s advantage is clear - it was able to finish computing the MCSs for all the reactions in under 14 seconds. Note that the dimensions stated in the tables are the ones before compression is applied. The conclusion is there are models for which it was not feasible to enumerate the full set of MCSs for a given target reaction before our work, but it is feasible now with MCS^2^.

**Table 1.** Result of running the methods on the hepatic polyamine and sulfur amino acid network [RPMSJM12]. *m* = 53*, n* = 73; target reaction 1.

| All times are<br>in seconds. | Optimized<br>Berge | Improved<br>MFK | Target-specific<br>Dual network | MCS<sup>2</sup><br>Dual network |
| --- | --- | --- | --- | --- |
| extreme ray computation | 270.2 | 270.2 | 1191.9 | 79.8 |
| secondary process time | >18000 | >18000 | 591.3 | 157.4 |
| total time | >18000 | >18000 | 1783.2 | 237.2 |

**Table 2.** Result of running the methods on the kinetic model of yeast network [SLS^+^13] with *m* = 295*, n* = 285; all the reactions were used as targets.

| All times are<br>in seconds. | Optimized<br>Berge | Improved<br>MFK | Target-specific<br>Dual network | MCS<sup>2</sup><br>Dual network |
| --- | --- | --- | --- | --- |
| extreme ray computation | 86.0 | 86.0 | >18000 | 53.0 |
| secondary process time | >18000* | >18000 | - | 13.6 |
| total time | >18000* | >18000 | >18000 | 66.6 |

\*: Berge computed the MCSs for the first five reactions before running out of time. The MFK and target-specific dual methods were not able to finish the computation of the MCSs even for the first reaction.

**Table 3.** Result of running the methods on Fernandez2006 ModelB [FSGLN06] with *m* = 75*, n* = 152; target reaction 1.

| All times are<br>n seconds. | Optimized<br>Berge | Improved<br>MFK | Target-specific<br>Dual network | MCS<sup>2</sup><br>Dual network |
| --- | --- | --- | --- | --- |
| extreme ray computation | 99.5 | 99.5 | >18000 | >18000 |
| secondary process | 2.1 | 1445.1 | - | - |
| total time | 101.6 | 1544.6 | >18000 | >18000 |

This is an example where MCS<sup>2</sup> and the target-specific dual methods could not finish in time, while the Berge and MFK methods reported all 194,689 MCSs for the compressed network’s reaction 1 fairly quickly.

We ran the MILP versions on larger networks alongside the MILP version of the target-specific dual approach, as described in [vKK14]. This version is also a part of CellNetAnalyzer and is believed to be the state-of-the-art for extracting some of the smallest MCSs in increasing order of size. We were able to compute 100 MCSs for reaction 10 (the first reaction with at least 100 MCSs) in the Li2012 Model [LSLN12], which has 578 reactions after compression. The time required by both approaches on the first 40 MCSs is shown in Figure 4.

**Figure 4:**
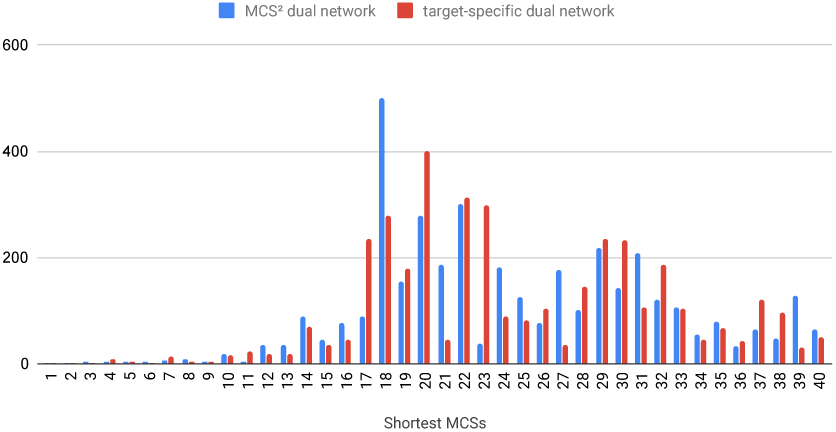
Time (in seconds) for computing each of the 40 smallest MCSs for reaction 10 (the first reaction which has at least 100 MCSs) of the Li2012 Calcium-mediated synaptic plasticity model [LSLN12]

Mixed-integer linear programming was used to find a subset of the MCSs [SGMR17]. The target-specific dual method has previously been used for this task, in a method called MCSEnumerator [vKK14, VMRR16]. As its authors state, not all the EFMs in the dual space result in valid MCSs, but by adding the appropriate constraints, one can remove the redundant results from the ILP’s feasible space. To get a sense of how our approach, MCS^2^-MILP, performs compared to MCSEnumerator, we implemented the MILP described in [vKK14], currently part of CellNetAnalyzer. We ran our implementation of MCSEnumerator [vKK14] and MCS^2^-MILP on *E.coli iAF1260* for the sake of comparison, which showed a similar performance, as shown in Table 4. This table contains the result of running MILPs for each reaction as a target reaction once per iteration. In each iteration we restricted the MILPs to not spend more than one minute on finding MCSs. Table 5

**Table 4.** Result of running MCS^2^-MILP and MCSEnumerator on the E coli iAF1260 network with 2382 reactions (981 reactions after compression).

| Method<br>used | Average Number<br>of MCSs | Average time for<br>shortest MCS | Number of targets<br>MILP failed on |
| --- | --- | --- | --- |
| MCS<sup>2</sup>-MILP | 12.74 MCSs | 4.45 seconds | 17 |
| MCSEnumerator | 12.07 MCSs | 5.22 seconds | 13 |

**Table 5.** Result of running MCS^2^-MILP and MCSEnumerator on the models from the BiGG database which initially have 2000 to 2600 reactions.

| Model ID | Average time for shortest MCS for MCSEnumerator | Average time for shortest MCS for MCS<sup>2</sup>-MILP | Reactions before (after) compression |
| --- | --- | --- | --- |
| iJO1366 | 4.66 seconds | 3.98 seconds | 2583 (1106) |
| iRC1080 | 7.12 seconds | 7.19 seconds | 2191 (1080) |
| STM_v1_0 | 1.82 seconds | 1.83 seconds | 2545 (1031) |
| iSbBS512_1146 | 14.10 seconds | 19.03 seconds | 2591 (1018) |
| iAF1260 | 5.22 seconds | 4.45 seconds | 2382 (981) |
| iSDY_1059 | 8.00 seconds | 9.63 seconds | 2539 (942) |
| iYL1228 | 1.88 seconds | 2.11 seconds | 2262 (805) |

## 5 Conclusions and Future Work

One key advantage of our method is that it does not depend on the target reaction to construct the dual network. The computations for one target reaction can therefore be reused for a different target reaction. Furthermore, it tends to operate on a smaller network than the original.

One limitation to our method is that it is primarily designed for single target reactions (rather than a target containing a set of reactions), while both are just as easily handled by the competitor methods. Although MCS^2^ does not find the MCSs for a set of reactions directly, it can easily find the MCSs for each reaction individually, then prune any supersets from the union of these MCSs. An alternate strategy for computing MCSs is via mixed-integer linear programming (MILP), particularly when only a few short sets are required, rather than a complete enumeration [RPdF^+^13, RPT^+^14]. We showed that MCS^2^ can be easily adapted to this task via the MCS^2^-MILP method, which has shown performance not inferior to that of the state-of-the-art.

Another strategy is to alter the double description method to directly find rays with minimal coordinated support instead of minimal support, e.g., by ignoring some of dimensions of the reconfigured network. Here it is important to be careful about zero-cycle flux modes, which are flux modes that have fluxes in both direction of a split reversible reaction. These are not valid flux modes, but they do appear in the output of the double description method [GK04] and they may cause the omission of some rays which contain them in their support.

As we mentioned, there are many models for which our method outperforms all other existing methods, while for some models, the best performance is obtained by the Berge algorithm. The challenge is to find out what features of these models are different, and then to decide ahead of time what method to choose for a given model.

Our method is based on novel insights, and may be refined further. Possible additional sources of improvement include identifying and removing unwanted supersets during the execution of the double description method and optimizing the process of superset removal during post-processing. We believe that our method opens the door to further ideas exploring this different kind of duality between EFMs and MCSs, and deeper insights into the structure of metabolic network models.

## References

[Ber84] Claude Berge. Hypergraphs: combinatorics of finite sets, volume 45 of North-Holland Mathematical Library. Elsevier, 1984.

[BMKP14] Aarash Bordbar, Jonathan Monk, Zachary King, and Bernhard Palsson. Constraint-based models predict metabolic and associated cellular functions. Nature Reviews Genetics, 15(2):107, 2014.

[BvKKH11] Kathrin Ballerstein, Axel von Kamp, Steffen Klamt, and Utz-Uwe Haus. Minimal cut sets in a metabolic network are elementary modes in a dual network. Bioinformatics, 28(3):381–387, 2011.

[Chi10] Leonid Chindelevitch. Extracting Information from Biological Networks. PhD thesis, MIT, 2010.

[CTRB14] Leonid Chindelevitch, Jason Trigg, Aviv Regev, and Bonnie Berger. An exact arithmetic toolbox for a consistent and reproducible structural analysis of metabolic network models. Nature Communications, 5, 2014.

[EMG08] Thomas Eiter, Kazuhisa Makino, and Georg Gottlob. Computational aspects of monotone dualization: A brief survey. Discrete Applied Mathematics, 156(11):2035–2049, 2008.

[FK96] Michael Fredman and Leonid Khachiyan. On the complexity of dualization of monotone disjunctive normal forms. Journal of Algorithms, 21(3):618–628, 1996.

[FSGLN06] Éric Fernandez, Renaud Schiappa, Jean-Antoine Girault, and Nicolas Le Novère. DARPP-32 is a robust integrator of dopamine and glutamate signals. PLoS Computational Biology, 2(12):e176, 2006.

[GDVL17] Andrew Gainer-Dewar and Paola Vera-Licona. The minimal hitting set generation problem: algorithms and computation. SIAM Journal on Discrete Mathematics, 31(1):63–100, 2017.

[GK04] Julien Gagneur and Steffen Klamt. Computation of elementary modes: a unifying framework and the new binary approach. BMC Bioinformatics, 5(1):175, 2004.

[HBK16] Björn-Johannes Harder, Katja Bettenbrock, and Steffen Klamt. Model-based metabolic engineering enables high yield itaconic acid production by Escherichia coli. Metabolic Engineering, 38:29–37, 2016.

[HFR+14] Hassan Hartman, David Fell, Sergio Rossell, Peter Jensen, Martin Woodward, Lotte Thorndahl, Lotte Jelsbak, John Olsen, Anu Raghunathan, Simon Daefler, and Mark Poolman. Identification of potential drug targets in Salmonella enterica sv. Typhimurium using metabolic modelling and experimental validation. Microbiology, 160(6):1252–1266, 2014.

[HKS08] Utz-Uwe Haus, Steffen Klamt, and Tamon Stephen. Computing knock-out strategies in metabolic networks. Journal of Computational Biology, 15(3):259–268, 2008.

[IB08] Marcin Imielinski and Calin Belta. Exploiting the pathway structure of metabolism to reveal high-order epistasis. BMC Systems Biology, 2(1):40, 2008.

[IBM] IBM. CPLEX optimizer.

[KG04] Steffen Klamt and Ernst Gilles. Minimal cut sets in biochemical reaction networks. Bioinformatics, 20(2):226–234, 2004.

[Kla06] Steffen Klamt. Generalized concept of minimal cut sets in biochemical networks. Biosystems, 83(2-3):233–247, 2006.

[KLD+15] Zachary King, Justin Lu, Andreas Dräger, Philip Miller, Stephen Federowicz, Joshua Lerman, Ali Ebrahim, Bernhard Palsson, and Nathan Lewis. BiGG Models: A platform for integrating, standardizing and sharing genome-scale models. Nucleic Acids Research, 44:D515–D522, 2015.

[KSRG07] dSteffen Klamt, Julio Saez-Rodriguez, and Ernst Gilles. Structural and functional analysis of cellular networks with CellNetAnalyzer. BMC Systems Biology, 1(1):2, 2007.

[LB07] Abdelhalim Larhlimi and Alexander Bockmayr. Minimal direction cuts in metabolic networks. In AIP Conference Proceedings, volume 940, pages 73–86. AIP, 2007.

[LDR+10] Chen Li, Marco Donizelli, Nicolas Rodriguez, Harish Dharuri, Lukas Endler, Vijayalakshmi Chelliah, Lu Li, Enuo He, Arnaud Henry, Melanie Stefan, et al. BioModels Database: An enhanced, curated and annotated resource for published quantitative kinetic models. BMC Systems Biology, 4(1):92, 2010.

[LNP12] Nathan Lewis, Harish Nagarajan, and Bernhard Palsson. Constraining the metabolic genotype–phenotype relationship using a phylogeny of in silico methods. Nature Reviews Microbiology, 10(4):291, 2012.

[LSLN12] Lu Li, Melanie Stefan, and Nicolas Le Novère. Calcium input frequency, duration and amplitude differentially modulate the relative activation of calcineurin and CaMKII. PloS One, 7(9):e43810, 2012.

[MR16] Stefan Müller and Georg Regensburger. Elementary vectors and conformal sums in polyhedral geometry and their relevance for metabolic pathway analysis. Frontiers in Genetics, 7:90, 2016.

[MvKK15] Radhakrishnan Mahadevan, Axel von Kamp, and Steffen Klamt. Genome-scale strain designs based on regulatory minimal cut sets. Bioinformatics, 31(17):2844–2851, 2015.

[PRP04] Nathan Price, Jennifer Reed, and Bernhard Palsson. Genome-scale models of microbial cells: evaluating the consequences of constraints. Nature Reviews Microbiology, 2(11):886, 2004.

[RPdF+13] Alberto Rezola, Jon Pey, Luis de Figueiredo, Adam Podhorski, Stefan Schuster, Angel Rubio, and Francisco Planes. Selection of human tissue-specific elementary flux modes using gene expression data. Bioinformatics, 29(16):2009–2016, 2013.

[RPMSJM12] Armando Reyes-Palomares, Raúl Montañez, Francisca Sánchez-Jiménez, and Miguel Medina. A combined model of hepatic polyamine and sulfur amino acid metabolism to analyze S-adenosyl methionine availability. Amino Acids, 42(2-3):597–610, 2012.

[RPT+14] Alberto Rezola, Jon Pey, Luis Tobalina, Ángel Rubio, John Beasley, and Francisco Planes. Advances in network-based metabolic pathway analysis and gene expression data integration. Briefings in Bioinformatics, 16(2):265–279, 2014.

[SFD00] Stefan Schuster, David Fell, and Thomas Dandekar. A general definition of metabolic pathways useful for systematic organization and analysis of complex metabolic networks. Nature Biotechnology, 18(3):326, 2000.

[SGMR17] Hyun-Seob Song, Noam Goldberg, Ashutosh Mahajan, and Doraiswami Ramkrishna. Sequential computation of elementary modes and minimal cut sets in genome-scale metabolic networks using alternate integer linear programming. Bioinformatics, 33(15):2345–2353, 2017.

[SLS+13] Natalie Stanford, Timo Lubitz, Kieran Smallbone, Edda Klipp, Pedro Mendes, and Wolfram Liebermeister. Systematic construction of kinetic models from genome-scale metabolic networks. PloS One, 8(11):e79195, 2013.

[SSC18] Nafiseh Sedaghat, Tamon Stephen, and Leonid Chindelevitch. Speeding up dualization in the Fredman-Khachiyan algorithm B. In Leibniz International Proceedings in Informatics, volume 103. Schloss Dagstuhl-Leibniz-Zentrum fuer Informatik, 2018.

[TCWS06] Cong T Trinh, Ross Carlson, Aaron Wlaschin, and Friedrich Srienc. Design, construction and performance of the most efficient biomass producing E. coli bacterium. Metabolic Engineering, 8(6):628–638, 2006.

[TS08] Marco Terzer and Jörg Stelling. Large-scale computation of elementary flux modes with bit pattern trees. Bioinformatics, 24(19):2229–2235, 2008.

[vKK14] Axel von Kamp and Steffen Klamt. Enumeration of smallest intervention strategies in genome-scale metabolic networks. PLoS Computational Biology, 10(1):e1003378, 2014.

[vKK17] Axel von Kamp and Steffen Klamt. Growth-coupled overproduction is feasible for almost all metabolites in five major production organisms. Nature Communications, 8:15956, 2017.

[vKWvD15] Jan Bert van Klinken and Ko Willems van Dijk. FluxModeCalculator: an efficient tool for large-scale flux mode computation. Bioinformatics, 32(8):1265–1266, 2015.

[VMRR16] Vítor Vieira, Paulo Maia, Isabel Rocha, and Miguel Rocha. Development of an integrated framework for minimal cut set enumeration in constraint-based models. In 10th International Conference on Practical Applications of Computational Biology & Bioinformatics, pages 193–201. Springer, 2016.

[Wag04] Clemens Wagner. Nullspace approach to determine the elementary modes of chemical reaction systems. The Journal of Physical Chemistry B, 108(7):2425–2431, 2004.

[ZSBC18] Hooman Zabeti, Tamon Stephen, Bonnie Berger, and Leonid Chindelevitch. A duality-based method for identifying elemental balance violations in metabolic network models. In Leibniz International Proceedings in Informatics, volume 113. Schloss Dagstuhl-Leibniz-Zentrum fuer Informatik, 2018.

